# Screening of plant growth-promoting rhizobacterial strains for the degradation and utilization of exogenous pectin as a soul carbon source

**DOI:** 10.1101/2021.01.05.425326

**Authors:** Mohammad K. Hassan

## Abstract

Universal primers for *gyrB* were used to screen 79 strains of *Bacillus* species. The genomic DNA from each strain was extracted, and the PCR product of *gyrB* was amplified and purified for gene sequencing. Gene sequences were edited, and aligned, and *Bacillus amyloliquefaciens* subsp. *plantarum* (Bap) strains identified based on the *gyrB* phylogenetic tree. Two primers (*exuT* and *uxuB*) were designed to detect pectin-associated transporter gene *exuT* and D-mannonate oxidoreductase gene *uxuB*. The Bap strains were then screened for the presence of the *exuT* and *uxuB* genes. These two pectin-utilizing genes were confirmed in 57 and 54 Bap strains by PCR amplification. Second, an *in vitro* tests were conducted to screen 59 Bap strains using pectin as a sole carbon source to confirm the pectate lyase and utilizing activity. Pectate Agar (PA) and Tris-Spizizen Salts (TSS) medium were used for the *in vitro* pectate lyase and degradation assays.

## 1. Introduction

Determining the differences of *Bacillus* species among closely related groups using 16S rRNA gene sequence is challenging because of their high sequence similarity (Wang, 2007). *Bacillus amyloliquefaciens* subsp. *amyloliquefaciens* and *Bacillus amyloliquefaciens* subsp. *plantarum* are two subspecies of Ba that cannot be distinguished by 16S rRNA gene sequence for their less DNA sequence dissimilarity. DNA gyrase subunit B and type II topoisomerase are referred to as *gyrB* that exerts a vital role in DNA replication is widely distributed in bacterial species and subspecies level (Huang, 1996). The *gyrB* gene provides accurate and faster results than 16S rRNA gene sequence to identify bacterial species and subspecies (Yamamoto et al., 1999). The *gyrB* gene sequences from different bacterial groups such as *Pseudomonas* (Yamamoto et al., 1999), *Acinetobacter* (Yamamoto et al., 1999), *Mycobacterium* (Kasai et al., 2000), *Bacillus thuringiesis* (La Duc et al., 2004), and *E*.*coli* (Fukushima et al., 2002) have been used for phylogenetic tree analysis. However, the 16S rRNA gene sequence is also helpful for the identification of bacterial species (Joung, 2002). Screening of *Bacillus* species using the *gyrB* gene sequence is an initial process to identify bacteria at the subspecies level.

The overall goal of this part of the research was to screen 79 *Bacillus* species for identification of Bap strains and the presence of pectin-utilizing genes (*exuT* and *uxuB*) identification in 59 Bap strains. The objectives of this study were i) to identify *Bacillus amyloliquefaciens* subsp. *plantarum* with *gyrB* primers (UP-1 & UP-2r), and ii) to identify *exuT* and *uxuB* genes with *exuT* and *uxuB* primers in the Bap strains.

## 2. Materials and methods

### 2.1. DNA extraction of *Bacillus sp*

A total of 79 *Bacillus* strains were collected and subcultured on TSA plates at 28°C for 24 hr. Profusely-grown bacteria were harvested for genomic DNA extraction using E.Z.N.A. ® DNA Isolation Kit and DNA concentration measured by NanoDrop UV-Vis Spectrophotometer. *Bacillus* strains were screened with primers for *gyrB, exuT* and *uxuB*.

### 2.2. PCR amplification, gel electrophoresis, and purification of PCR product of *gyrB* gene

Master Mix (2X) was prepared by several components such as *Tag* DNA polymerase, dNTPS, and Mgcl_2_. Other components such as forward primers, reverse primers, DNA template and nuclease-free water were also added for Master Mix preparation. Touchdown PCR was used to avoid nonspecific sequence amplification, and UP-1 and UP-2r *gyrB* universal primers were used (Yamada et al., 1999).The first step of Touchdown PCR was a separation of DNA double strand through the heating process at 95°C called denaturation for 30 seconds. The second step was lowering the temperature at 65°C to allow primers to anneal to complementary sequences for 30 seconds, and the third step was to synthesize of DNA strand through DNA polymerase enzymes to create double strand called primer extension at 72°C for 40 seconds. These steps were repeated 15 times. In the second step, the annealing temperature was changed to 50°C. The remaining temperatures were the same and were repeated 30 times. The final stage was a primer extension at 72°C for 5 minutes. Eppendorf Thermal Mastercycler (Eppendorf AG, Hamburg) was used for Touchdown PCR cycles. Agarose gel (1%) and 1X Tris Acetate EDTA (TAE) buffer were used for gel electrophoresis. The voltage was 110V for 35 minutes. The Agarose gel was stained in Ethidium bromide for 10 minutes. And was then properly de-stained for 5 minutes to remove ethidium bromide. The gel images were photographed using the AlphaImager® HP high-performance imaging System. PCR product was purified by E.Z.N.A. Cycle Pure Kit. The cleaned PCR reactions (5 µl) were taken from 40 µl PCR reaction. One µl of the forward and the reverse primers from 20 µM primers were separately added to 5 µl of the cleaned PCR reaction, The total PCR reactions were 6 µl for forward and 6 µl for reverse primers. The cleaned PCR reaction samples were sent to Lucigen (USA) for sequencing. Genomic sequences were edited and analyzed by Chromas Pro software in FASTA format and aligned by CLC Genomic Workbench software.

### 2.3. *gyrB* phylogenetic tree construction

MEGA6 software was used for *gyrB* phylogenetic tree construction. ClustalW alignment and Maximum likelihood statistical method were used for gene sequence alignment and phylogenetic tree. The Bap and other *Bacillus sp*. sequences were collected from Genbank and aligned with 76 strains to compare among the closely related species and subspecies. Bap sequences of gene bank and collected strains were compared for their similarities and dissimilarities.

### 2.4. *exuT* and *uxuB* primer design

Hexuronate transporter (*exuT*) primers 211F and 1070R were collected from, Dr. Mark Liles lab. D-mannonate/D-fructuronate oxidoreductase (*uxuB*) Primers 60F and 696R were designed for identification of Bap strains. Two-gene sequences (*exuT* and *uxuB*) of various Bap were collected from NCBI Genbank for primer design. Primer designer V. 2.0 software was used for *uxuB* primers. After designing the *uxuB* primers, the primers’parameters were checked by the online-based prediction tools sequence manipulation suite software interface (P, 2000).

### 2.5. PCR amplification and gel electrophoresis of *exuT* and *uxuB* genes

The touchdown PCR method was used to avoid nonspecific sequence amplification. The parameters for touchdown PCR conditions were same as a *gyrB* gene sequence.

## 3. Results

### 3.1. DNA extraction of *Bacillus sp*

**Table 1.**
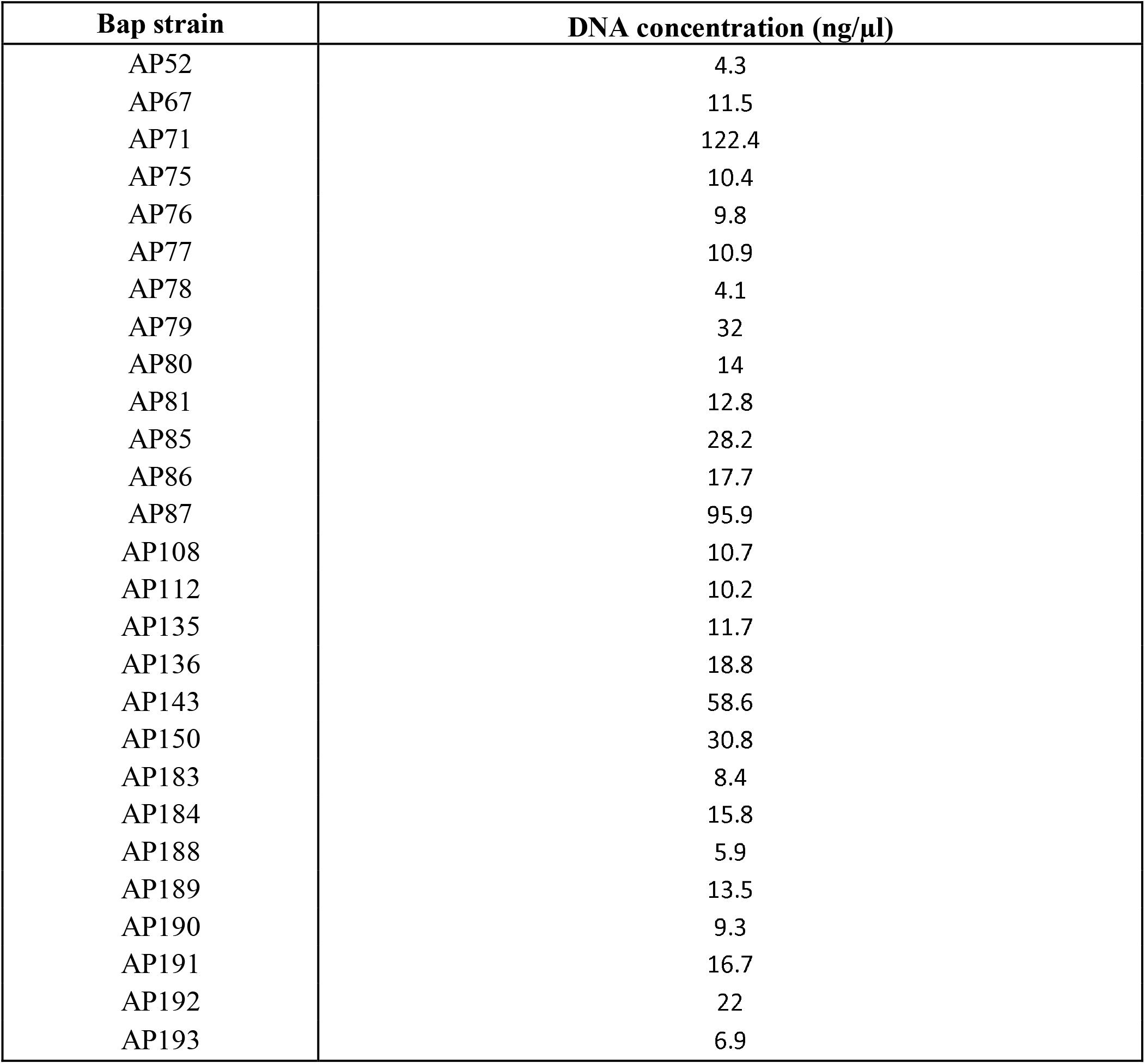
DNA concentration of 27 Bap strains.

**Table 2.**
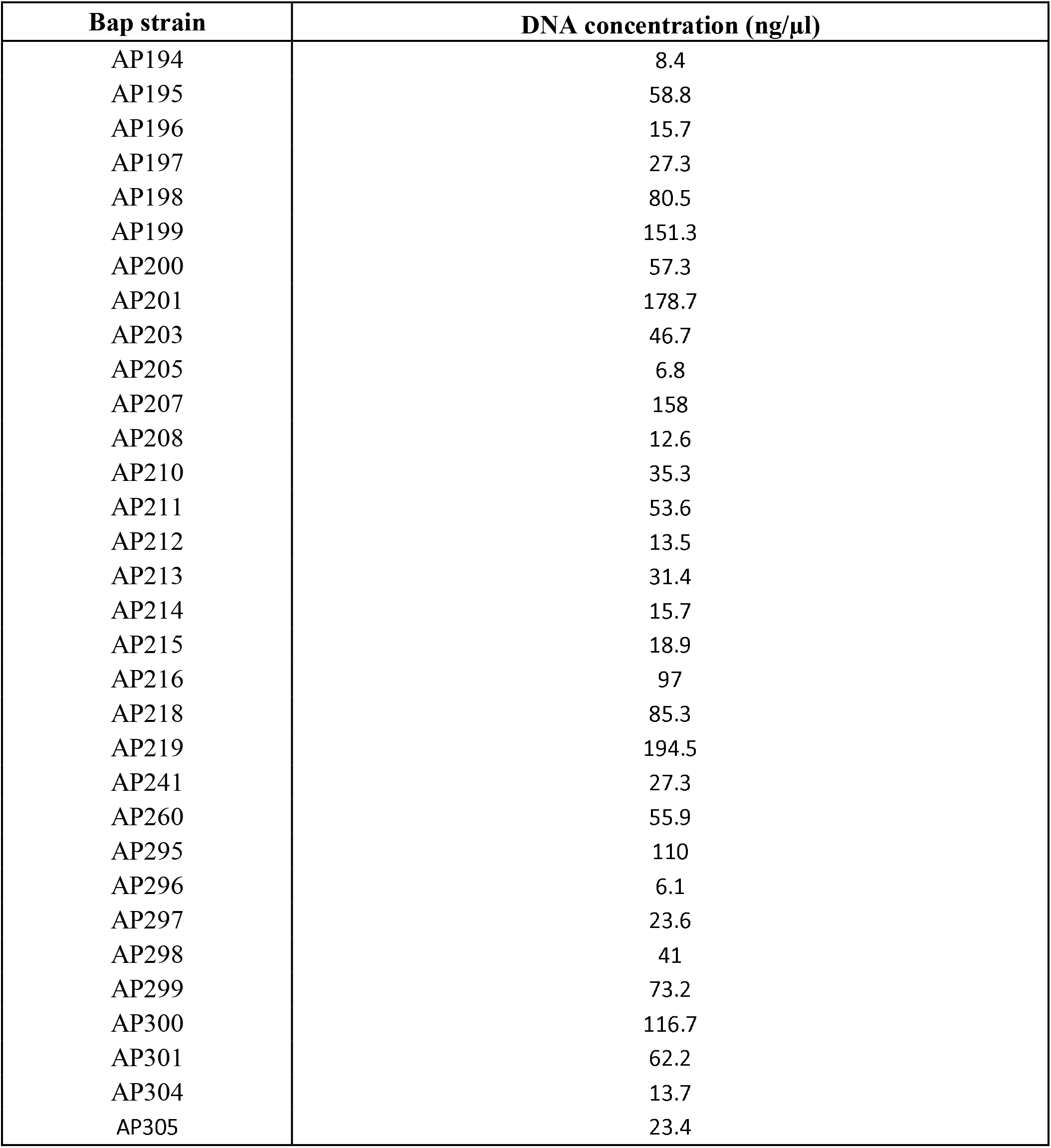
DNA concentration of 33 Bap strains.

| Bap strain | DNA concentration (ng/μl) |
| --- | --- |
| AP194 | 8.4 |
| AP195 | 58.8 |
| AP196 | 15.7 |
| AP197 | 27.3 |
| AP198 | 80.5 |
| AP199 | 151.3 |
| AP200 | 57.3 |
| AP201 | 178.7 |
| AP203 | 46.7 |
| AP205 | 6.8 |
| AP207 | 158 |
| AP208 | 12.6 |
| AP210 | 35.3 |
| AP211 | 53.6 |
| AP212 | 13.5 |
| AP213 | 31.4 |
| AP214 | 15.7 |
| AP215 | 18.9 |
| AP216 | 97 |
| AP218 | 85.3 |
| AP219 | 194.5 |
| AP241 | 27.3 |
| AP260 | 55.9 |
| AP295 | 110 |
| AP296 | 6.1 |
| AP297 | 23.6 |
| AP298 | 41 |
| AP299 | 73.2 |
| AP300 | 116.7 |
| AP301 | 62.2 |
| AP304 | 13.7 |
| AP305 | 23.4 |

The highest DNA concentration was 194.5 ng/µl in Bap strain AP218, and the lowest DNA concentration was 4.3 ng/µl in Bap strain AP52. The average DNA concentration was 44.21 ng/µl.

### 3.2. PCR, Gel electrophoresis, and Purification of PCR product of *gyrB* gene

**Table 3.**
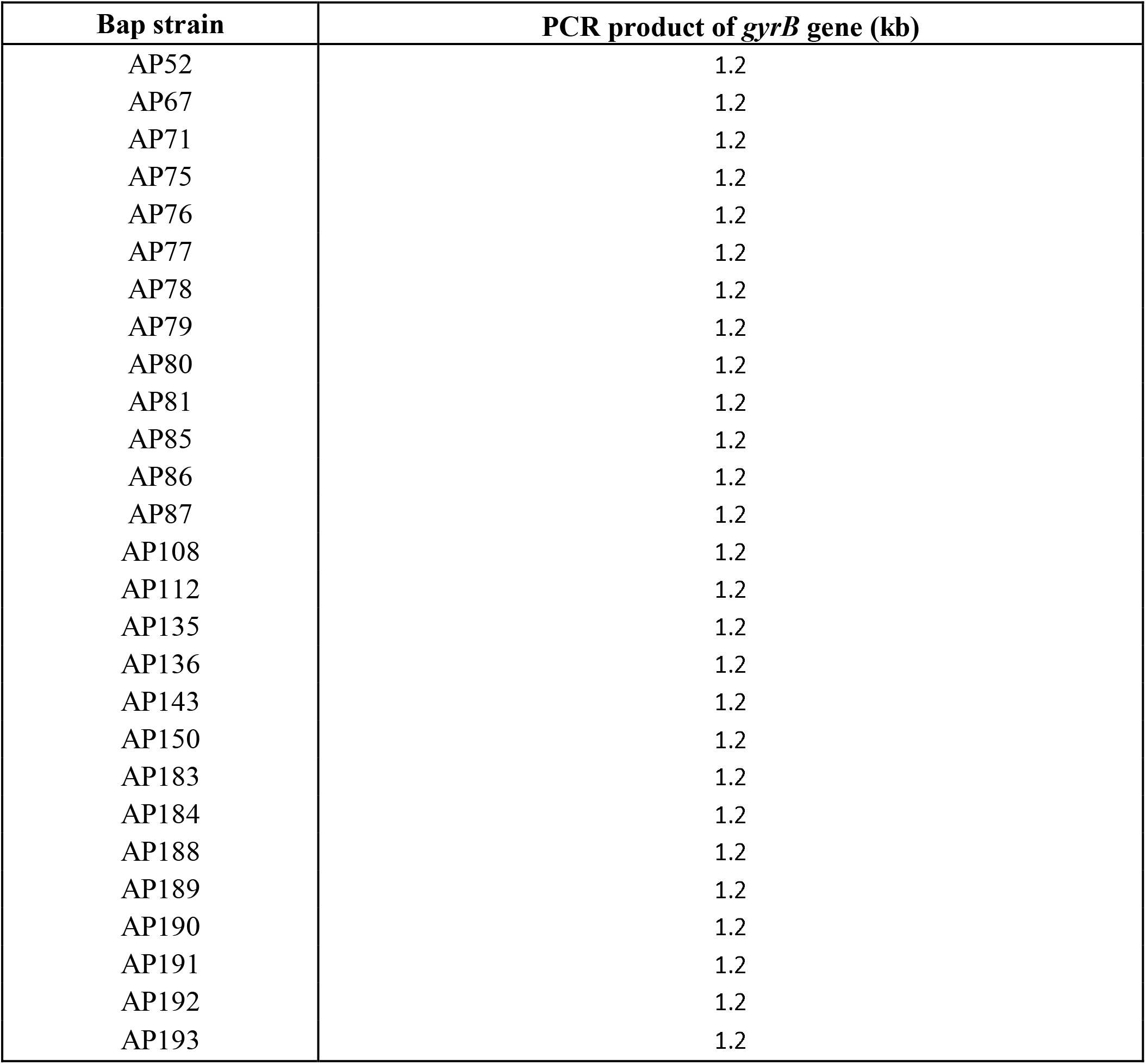
PCR product size of *gyrB* gene (kb) in 27 *Bacillus* sp.

| Bap strain | PCR product of *gyrB* gene (kb) |
| --- | --- |
| AP52 | 1.2 |
| AP67 | 1.2 |
| AP71 | 1.2 |
| AP75 | 1.2 |
| AP76 | 1.2 |
| AP77 | 1.2 |
| AP78 | 1.2 |
| AP79 | 1.2 |
| AP80 | 1.2 |
| AP81 | 1.2 |
| AP85 | 1.2 |
| AP86 | 1.2 |
| AP87 | 1.2 |
| AP108 | 1.2 |
| AP112 | 1.2 |
| AP135 | 1.2 |
| AP136 | 1.2 |
| AP143 | 1.2 |
| AP150 | 1.2 |
| AP183 | 1.2 |
| AP184 | 1.2 |
| AP188 | 1.2 |
| AP189 | 1.2 |
| AP190 | 1.2 |
| AP191 | 1.2 |
| AP192 | 1.2 |
| AP193 | 1.2 |

**Table 4.** PCR product size of *gyrB* gene (kb) in 33 *Bacillus* sp.

| Bap strain | PCR product of *gyrB* gene (kb) |
| --- | --- |
| AP194 | 1.2 |
| AP195 | 1.2 |
| AP196 | 1.2 |
| AP197 | 1.2 |
| AP198 | 1.2 |
| AP199 | 1.2 |
| AP200 | 1.2 |
| AP201 | 1.2 |
| AP203 | 1.2 |
| AP205 | 1.2 |
| AP207 | 1.2 |
| AP208 | 1.2 |
| AP210 | 1.2 |
| AP211 | 1.2 |
| AP212 | 1.2 |
| AP213 | 1.2 |
| AP214 | 1.2 |
| AP215 | 1.2 |
| AP216 | 1.2 |
| AP218 | 1.2 |
| AP219 | 1.2 |
| AP241 | 1.2 |
| AP260 | 1.2 |
| AP295 | 1.2 |
| AP296 | 1.2 |
| AP297 | 1.2 |
| AP298 | 1.2 |
| AP299 | 1.2 |
| AP300 | 1.2 |
| AP301 | 1.2 |
| AP304 | 1.2 |
| AP305 | 1.2 |

### 3.3. *gyrB* phylogenetic tree construction

Out of 79 *Bacillus* strains, 59 strains were identified as Bap strains. The remaining of the 19 bacterial strains were identified *B*.*mojavensis, B*.*sonorensis, B*.*tequilensis, Bacillus subtilis*, and *B*.*pumilis*. None of the strains were found to cluster together with *Bacillus amyloliquefaciens* subsp. *amyloliquefaciens*.

### 3.4. *exuT* and *uxuB* primer design

The best-predicted primers were GTTCTCTGTTCAGCAATG (60F), GAAAAGGCAAAAGACGAG (222F), CTTATTGGATCGGTCTGT (671R), and GTTTCATCGGTGTATGTG (696R). The 60F and 696R primers were selected for Bap strains screening. The sequence length of forward and reverse primers was 18. GC content (%) was 44.44, molecular weight (Daltons) was 5480.63, and nearest neighbor Tm (degrees C) was 56.15 in 60F *uxuB* primers. GC content (%) was 44.44, Molecular weight (Daltons) was 5445.62, and nearest neighbor Tm (degrees C) was 56.21 in 696R *uxuB* primers.

### 3.5. PCR amplification and gel electrophoresis of *exuT* and *uxuB* genes

**Table 5.** PCR product of *exuT* and *uxuB* gene (bp) in 27 Bap strains.

| Bap strain | PCR product of *exuT* gene (bp) | PCR product of *uxuB* gene (bp) |
| --- | --- | --- |
| AP52 | 900 | 600 |
| AP67 | 900 | 600 |
| AP71 | 900 | 600 |
| AP75 | 900 | 600 |
| AP76 | 900 | 600 |
| AP77 | 900 | 600 |
| AP78 | 900 | 600 |
| AP79 | 900 | 600 |
| AP80 | 900 | 600 |
| AP81 | 900 | 600 |
| AP85 | 900 | 600 |
| AP86 | 900 | 600 |
| AP87 | 900 | 600 |
| AP108 | 900 | 600 |
| AP112 | 900 | 600 |
| AP135 | 900 | 600 |
| AP136 | 900 | 600 |
| AP143 | 900 | 600 |
| AP150 | 900 | 600 |
| AP183 | 900 | 600 |
| AP184 | 900 | 600 |
| AP188 | 900 | 600 |
| AP189 | 900 | 600 |
| AP190 | 900 | 600 |
| AP191 | 900 | 600 |
| AP192 | 900 | 600 |
| AP193 | 900 | 600 |

**Table 6.** PCR product of *exuT* and *uxuB* gene (bp) in 30 and 25 Bap strains.

| Bap strain | PCR product of *exuT* gene (bp) | PCR product of *uxuB* gene (bp) |
| --- | --- | --- |
| AP194 | 900 | 600 |
| AP195 | 900 | 600 |
| AP196 | 900 | 600 |
| AP197 | 900 | 600 |
| AP198 | 900 | 600 |
| AP199 | 900 | 600 |
| AP200 | 900 | 600 |
| AP201 | 900 | 600 |
| AP203 | 900 | 600 |
| AP205 | 900 | 600 |
| AP207 | 900 | 600 |
| AP208 | 900 | 600 |
| AP211 | 900 | - |
| AP213 | 900 | 600 |
| AP214 | 900 | - |
| AP215 | 900 | 600 |
| AP216 | 900 | 600 |
| AP218 | 900 | - |
| AP219 | 900 | - |
| AP241 | 900 | 600 |
| AP260 | 900 | 600 |
| AP295 | 900 | 600 |
| AP296 | 900 | 600 |
| AP297 | 900 | 600 |
| AP298 | 900 | 600 |
| AP299 | 900 | 600 |
| AP300 | 900 | 600 |
| AP301 | 900 | - |
| AP304 | 900 | 600 |
| AP305 | 900 | 600 |

## 4. Discussion

The results from the *Bacillus* species genomic DNA extraction indicate that some of the strains have higher DNA concentration than other strains. Out of 79 *Bacillus* species, 16 had high DNA concentrations, and 30 had moderate DNA concentrations. The remaining strains showed low DNA concentrations. The length of the amplified *gyrB* gene had detected ∼1.2kb, and previous studies have reported the same size in *Bacillus* species (Wang et al., 2007).

The length of *gyrB* PCR product results infers that all the strains were *bacillus* species. None of the strains were found different bacterial species. In this way, eight different *Bacillus* species were identified based on *gyrB* PCR product amplification. Out of 59 Bap strains, *exuT* gene was identified in 56 Bap strains. A total of 59 Bap strains, *uxuB* gene was detected in 54 Bap strains. Three Bap strains (AP102, AP204, and AP214) were found *exuT and uxuB* gene negative.

The *gyrB* phylogenetic tree results revealed that Ba DSM7, Ba LL3, TA208, and Ba XH7 *gyrB* gene sequences collected from NCBI database are clustered as a single clade separated from the Bap strains. This single clade indicated that collected four strains were distinct subspecies of Ba, and none of the *Bacillus* species strains were associated with them. A total of 59 strains were grouped together with the reference strains indicating that they were Bap strains with high bootstrap values. A bootstrap replication value was used for constructing the *gyrB* phylogenetic tree. Bootstrap uses for making inferences and robustness, similar to the use of p values in statistical analysis (Holmes, 2003). Bootstrap value 95% or higher for a given branch indicates topology correctness at the branch (Nei, 2000). AP 82 and *B. cereus* were sister groups that indicate closest relatives in 99% bootstrap replications. *B. pumilus* and AP 70 were sister groups, and AP 100 was the outgroup in it. This type of relationship indicates monophyletic clade. Monophyletic clade consists of ancestor and descendants. Similarly, strong bootstrap support was found for *B. subtilus, B. mojavensis*, and *B. tequilensis. B. mojavensis* and AP242 were sister group with 99% bootstrap value. In this way, 79 *Bacillus* species were compared among reference strains collected from NCBI database.

---

**Figure 1:**
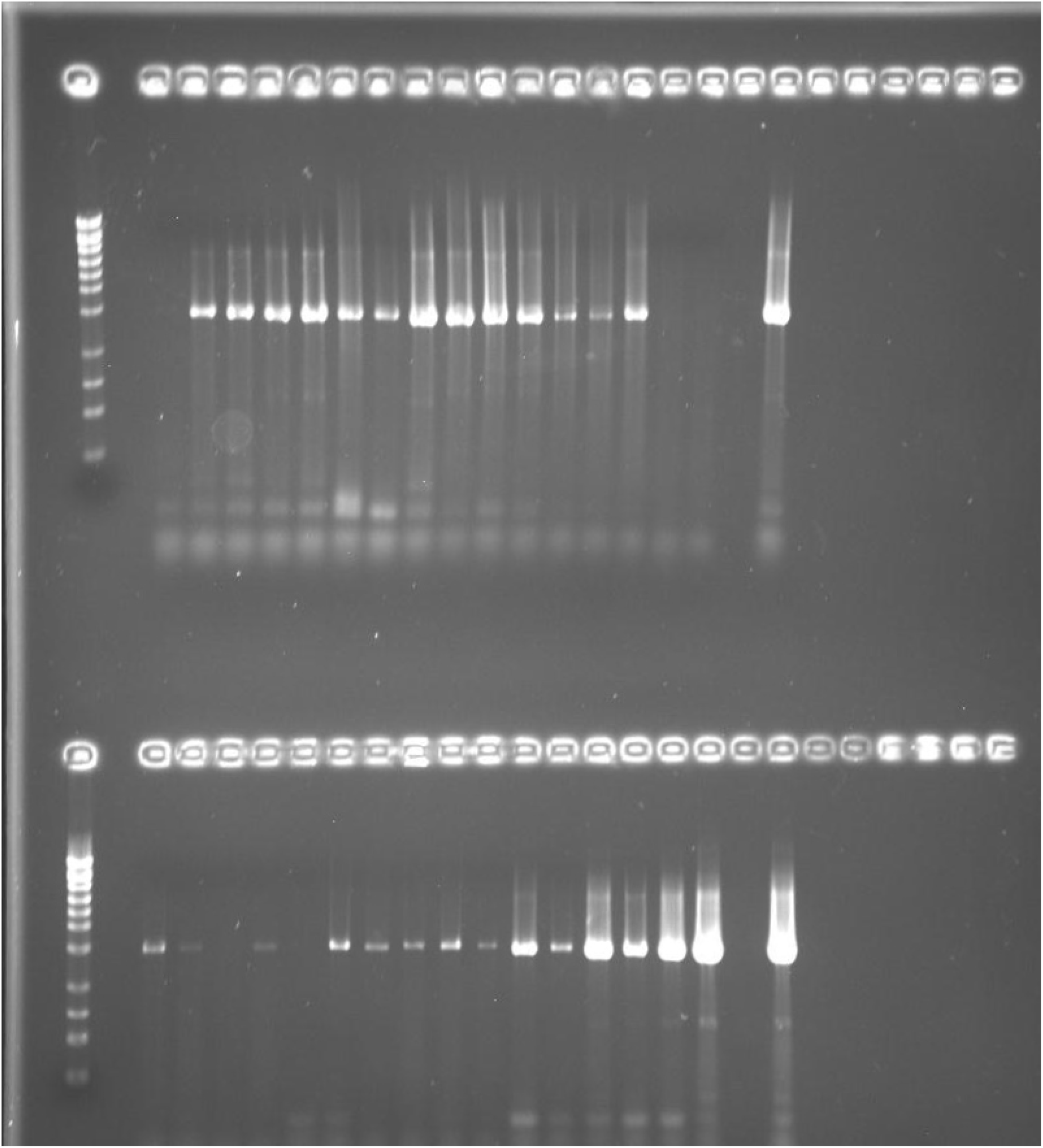
*gyrB* gene detected in *Bacillus* species (AP60, AP67, AP70, AP71, AP72, AP75, AP76, AP77, AP78, AP79, AP80, AP81, AP82, AP100, AP111, AP112, AP126, AP135, AP143, AP150, AP151, AP165, AP183, and AP188)

**Figure 2:**
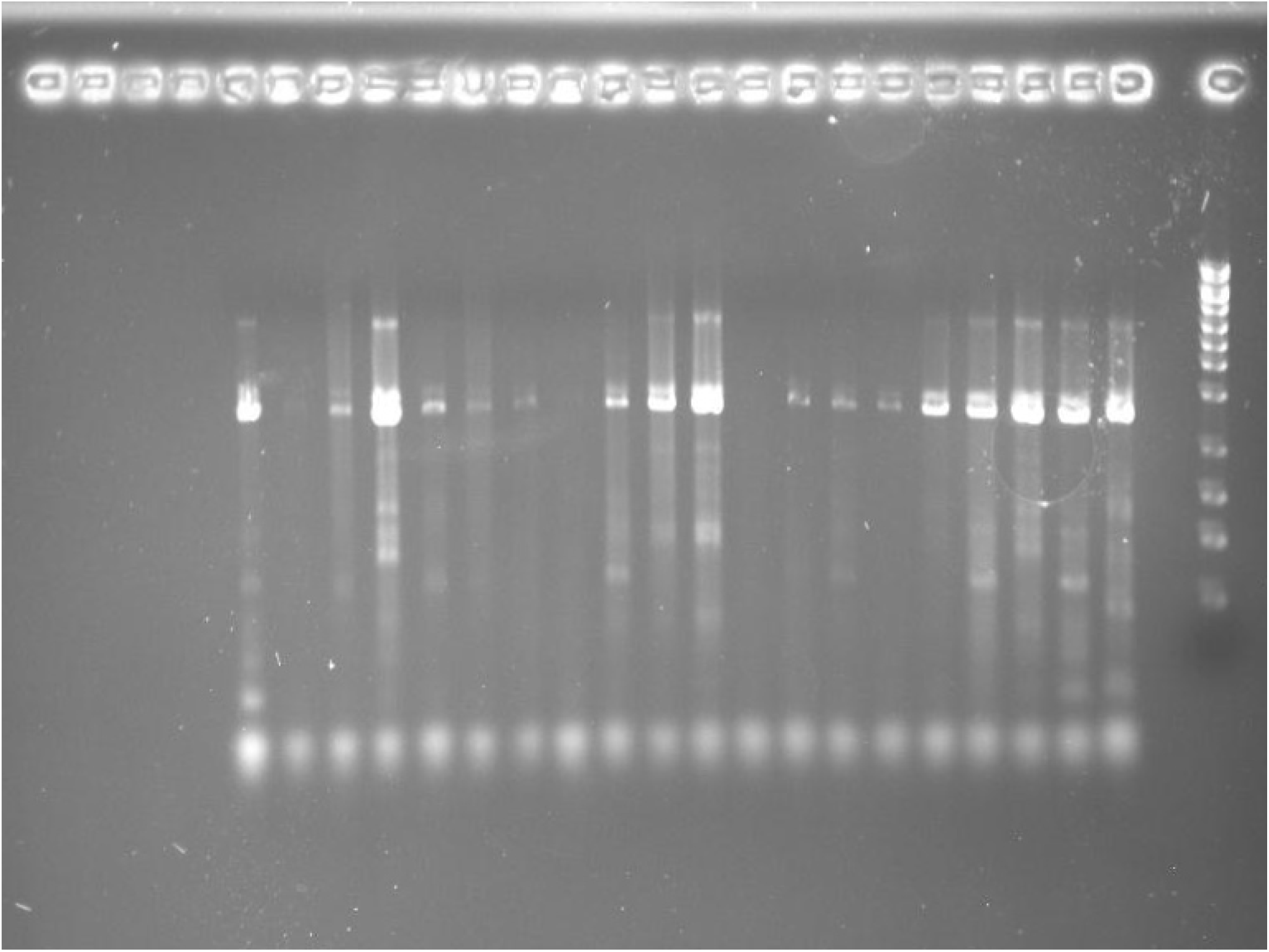
*gyrB* gene detected in *Bacillus* species (AP189, AP192, AP197, AP199, AP205, AP206, AP207, AP208, and AP210)

**Figure 3:**
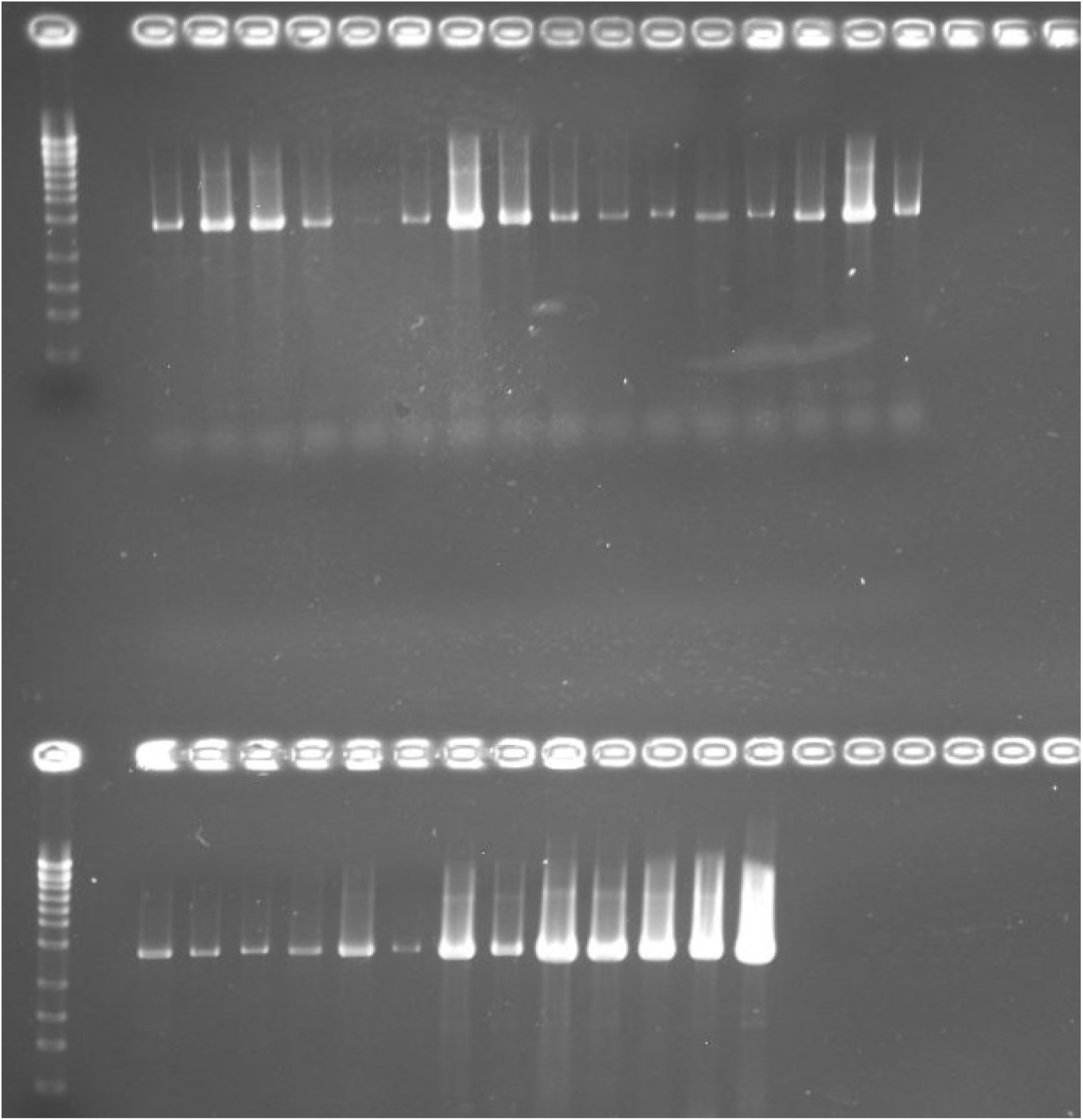
*gyrB* gene detected in *Bacillus* species (AP211, AP212, AP213, AP214, AP216, AP218, AP219, AP241, AP242, AP260, AP278, AP279, AP295, AP296, AP297, AP298, AP299, AP300, AP301, AP304, AP52, AP85, AP86, AP87, AP105, AP106, and AP108)

**Figure 4:**
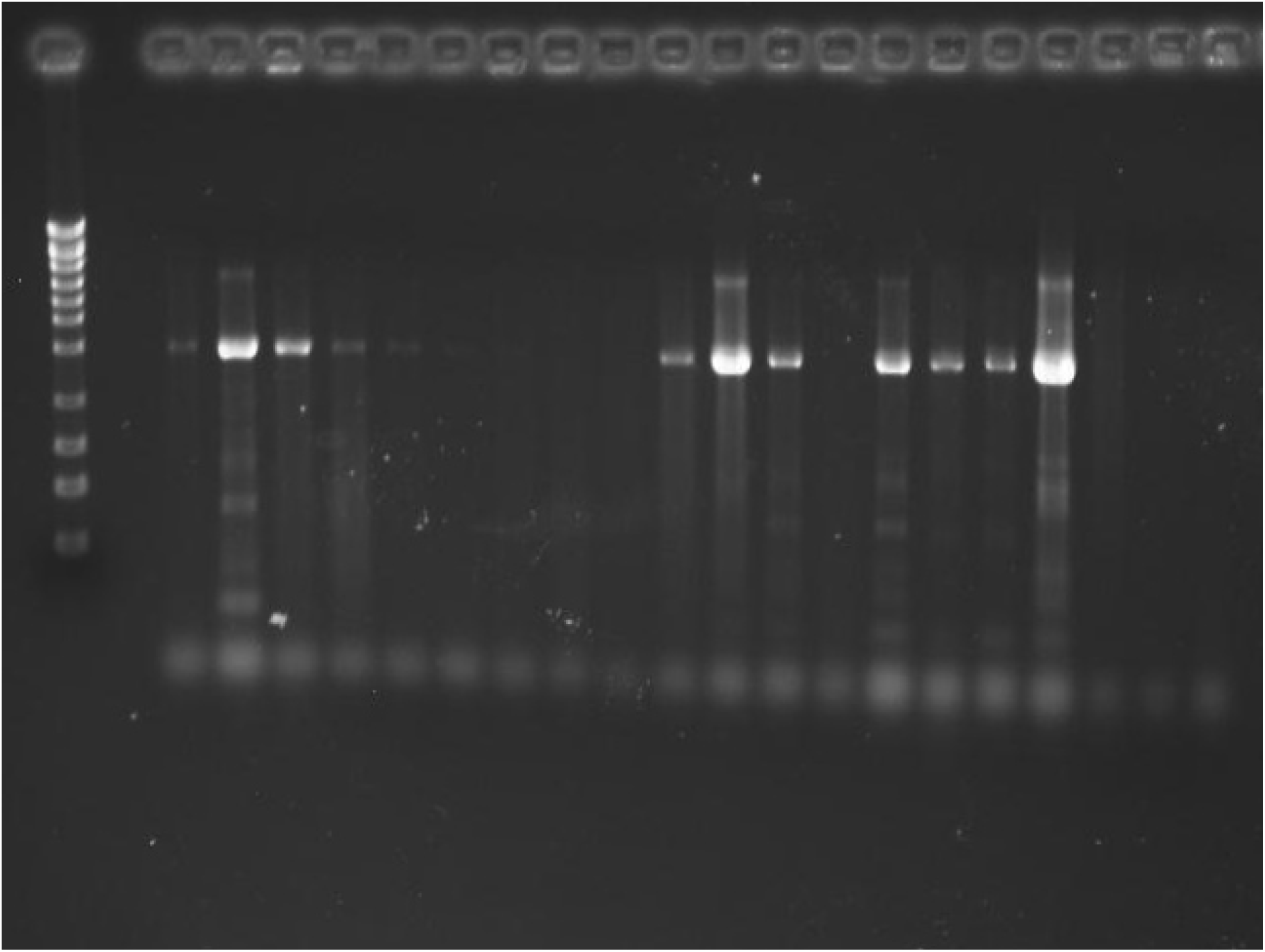
*gyrB* gene detected in *Bacillus* species (AP184, AP190, AP204, AP215, AP193, AP194, AP195, and AP196)

**Figure 5:**
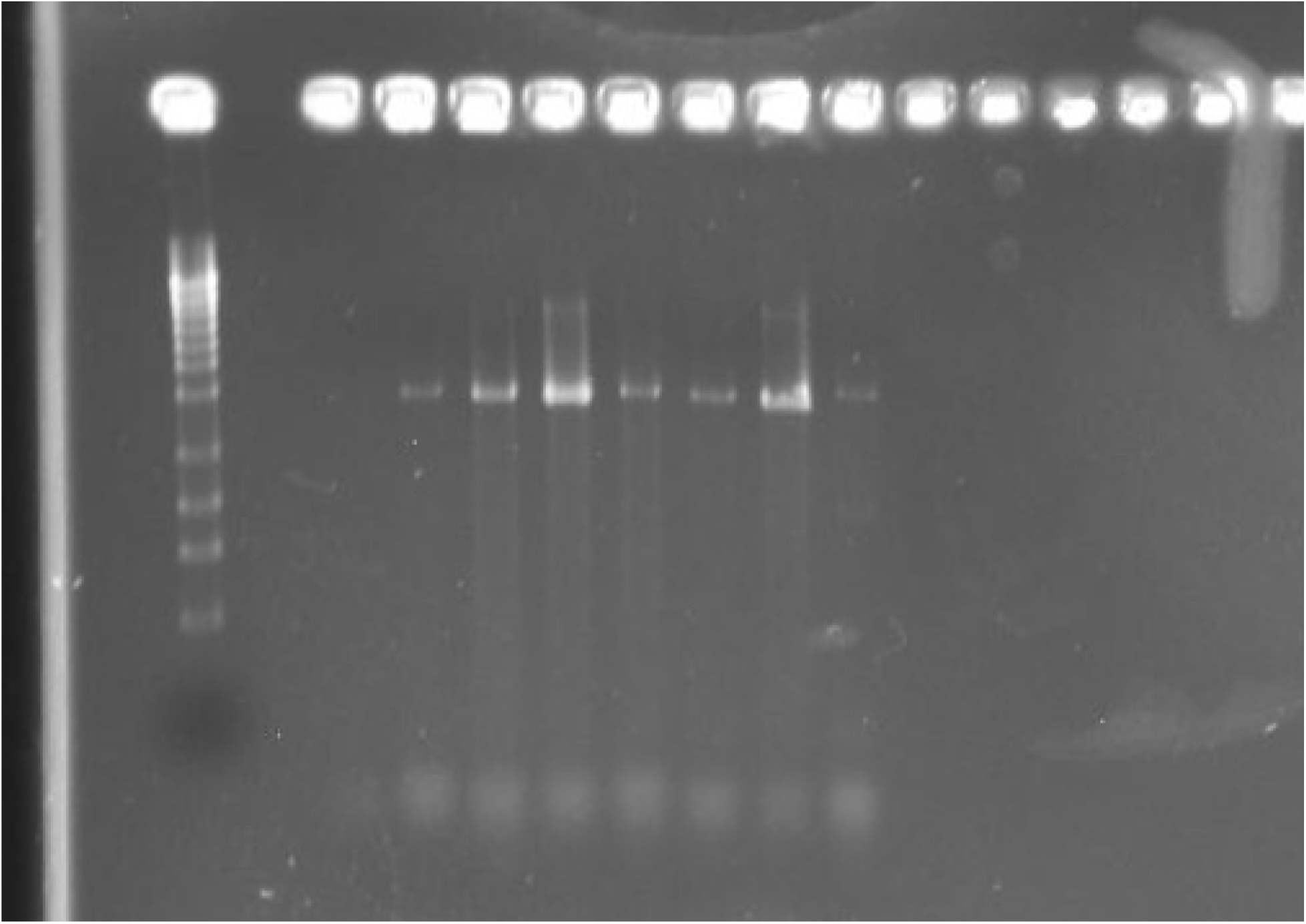
*gyrB* gene detected in *Bacillus* species (AP109, AP191, AP200, and AP200)

**Figure 6:**
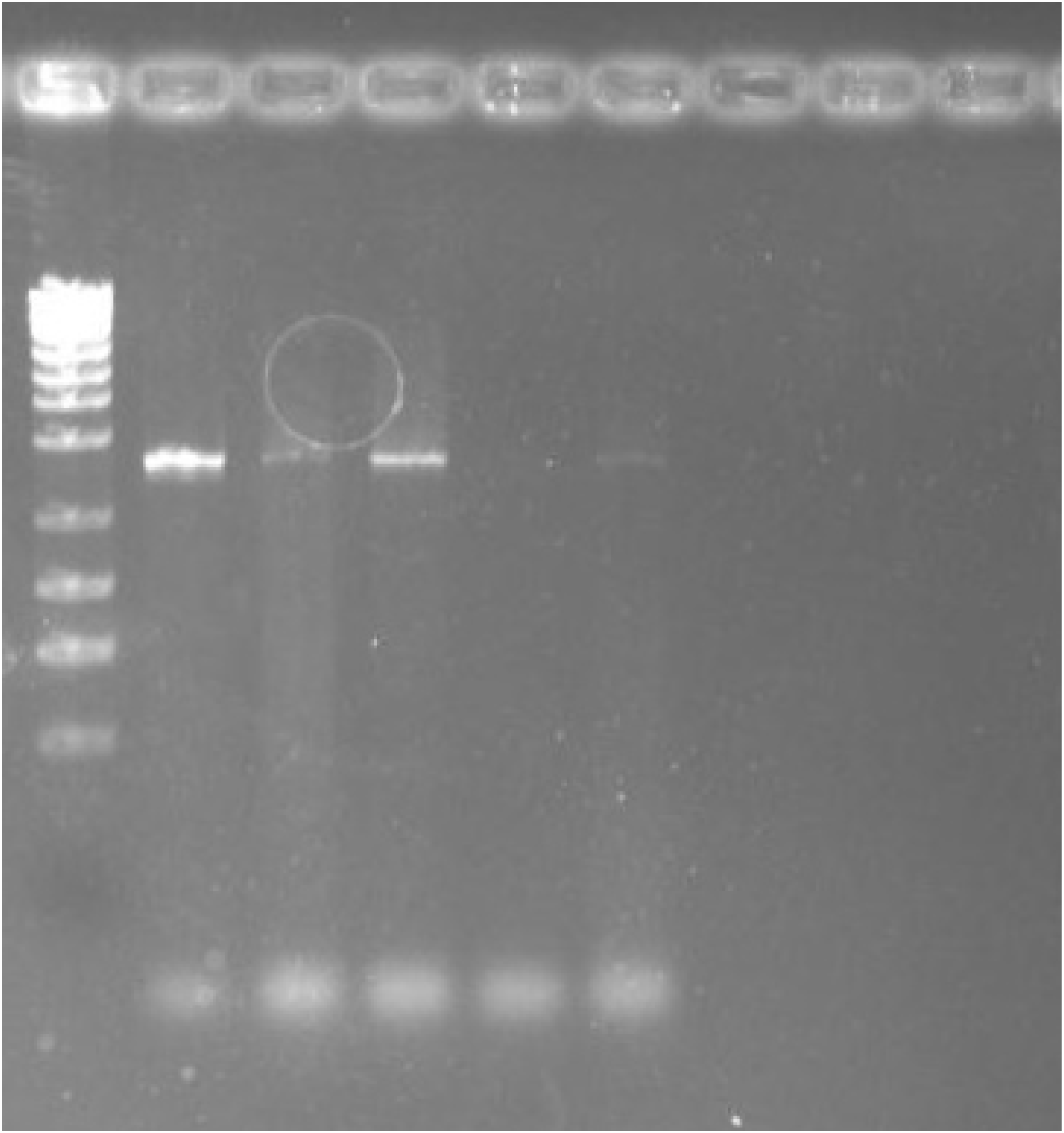
*gyrB* gene identified in *Bacillus* species (AP201 and AP204)

**Figure 7:**
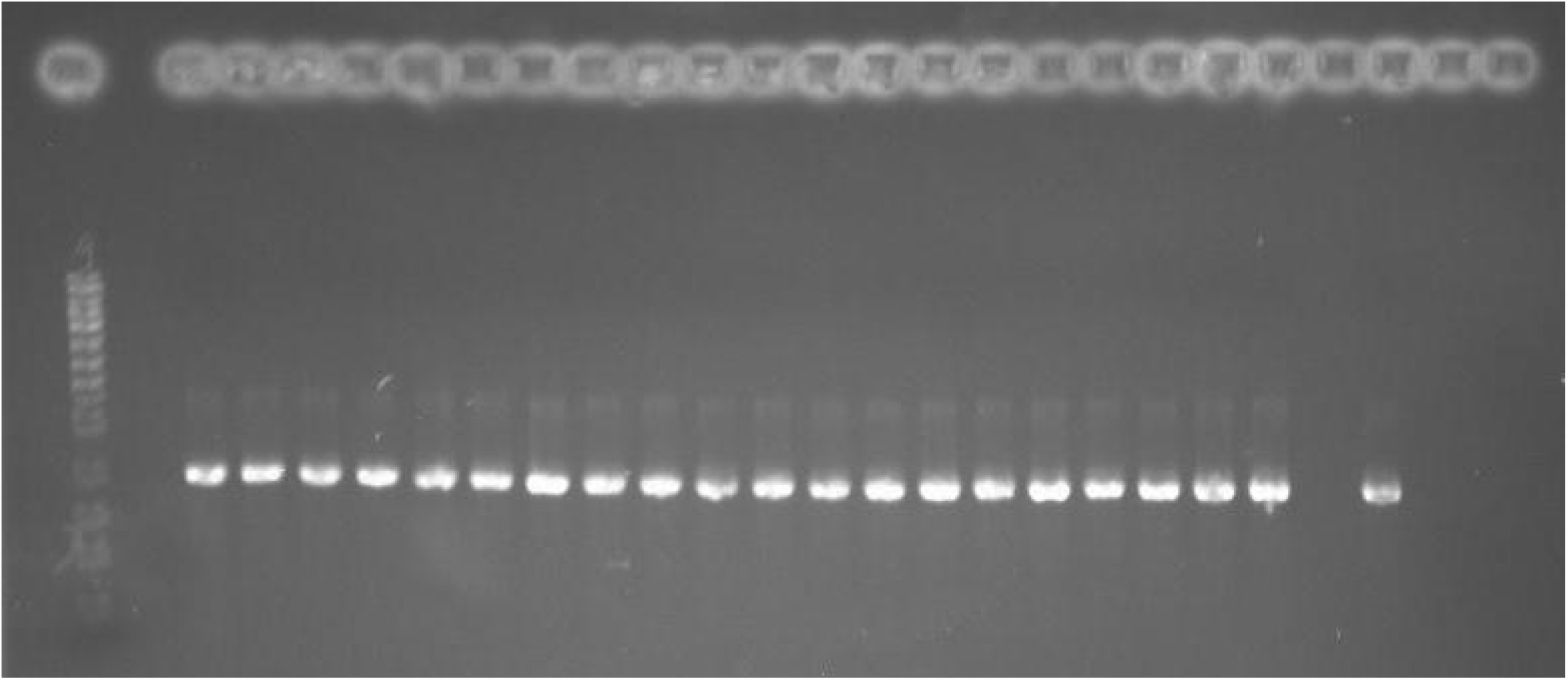
*exuT* gene detected in Bap strains (AP191, AP199, AP218, AP201, AP71, AP195, AP200, AP87, AP215, AP212, AP183, AP136, AP210, AP203, AP79, AP77, AP108, AP188, AP213, AP189, and AP193)

**Figure 8:**
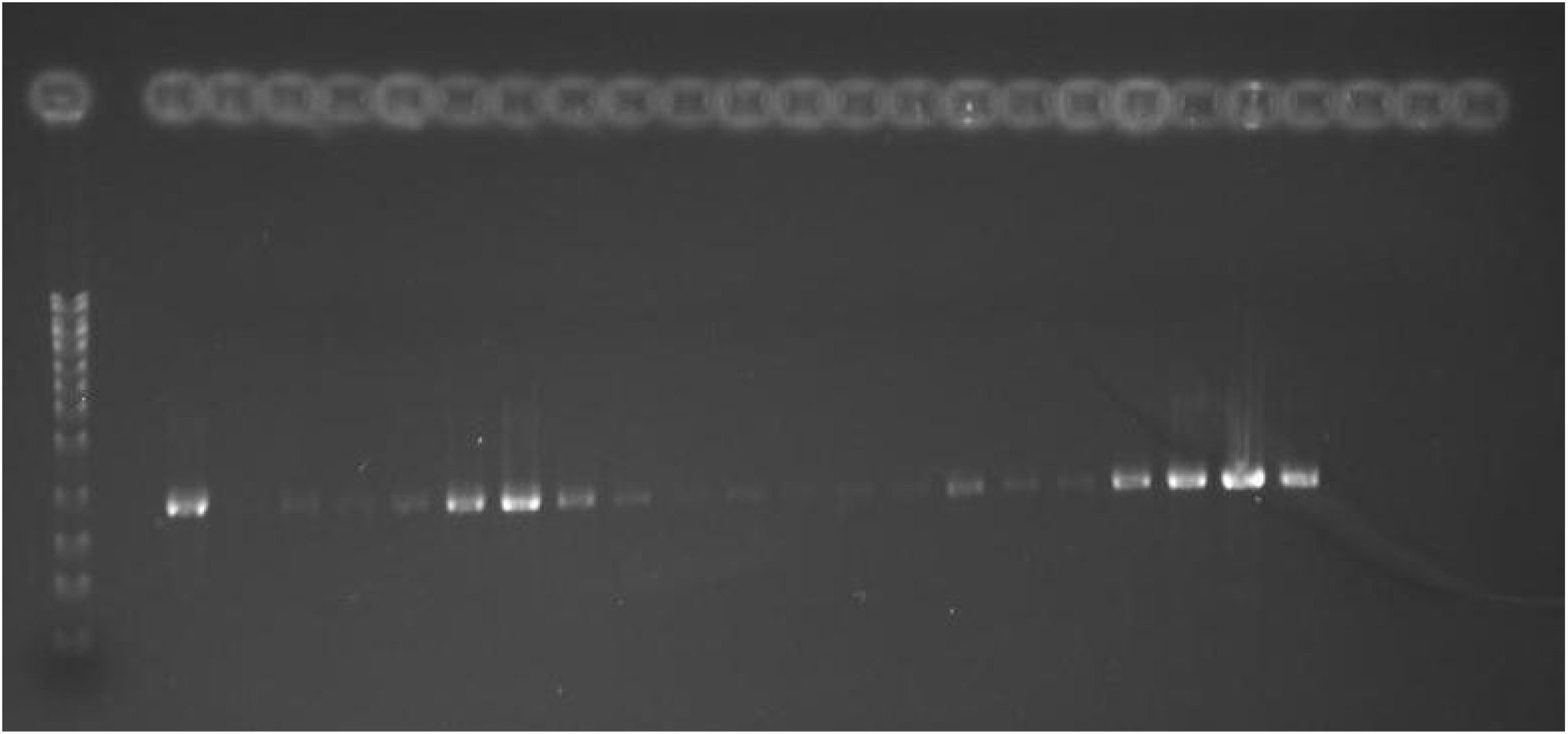
*exuT* gene detected in Bap strains (AP298, AP197, AP194, AP67, AP85, AP112, AP300, AP260, AP304, AP52, AP198, AP190, AP135, AP76, AP81, AP297, AP208, AP80, and AP193)

**Figure 9:**
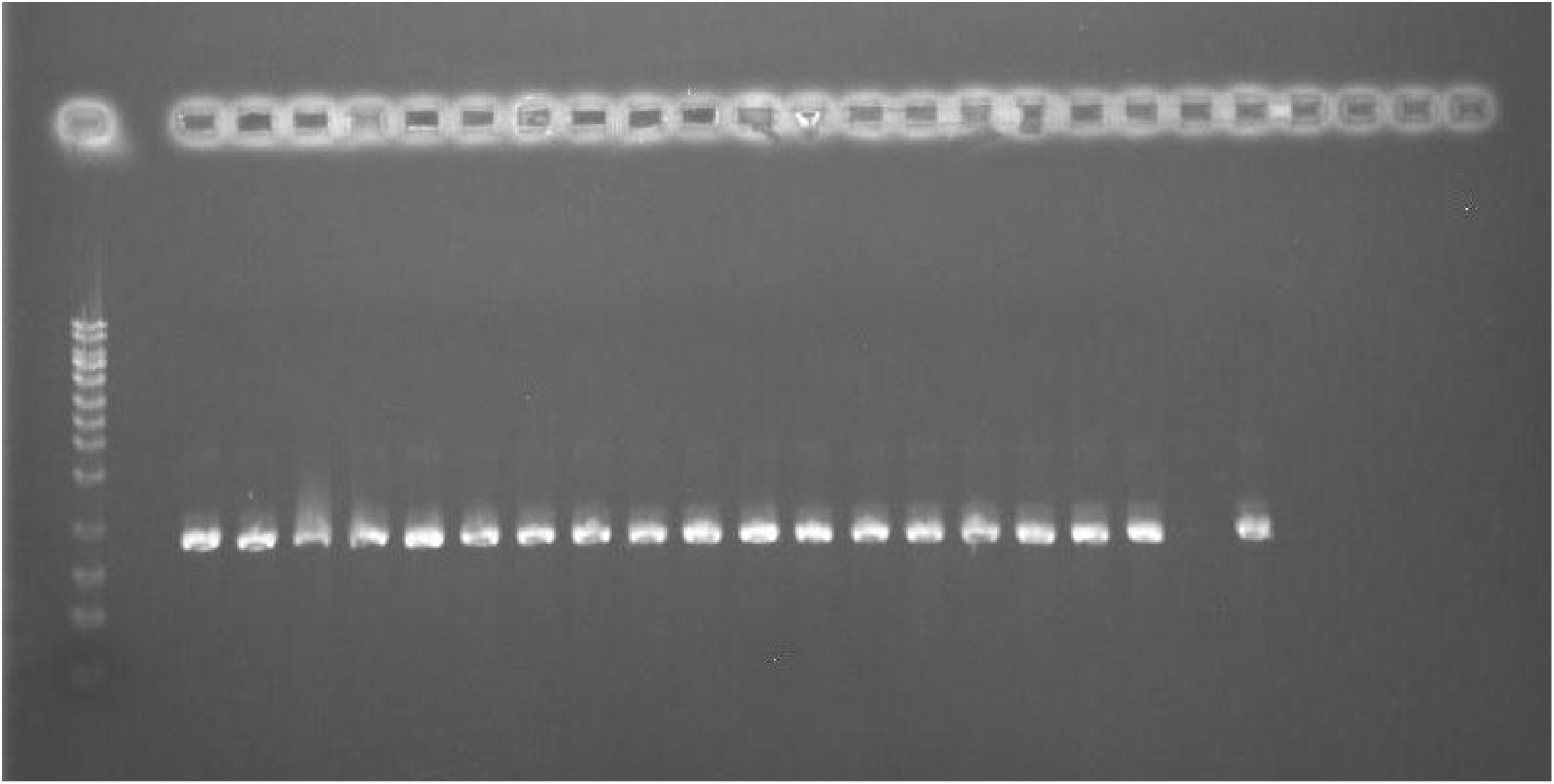
*exuT* gene detected in Bap strains (AP86, AP196, AP78, AP305, AP192, AP184, AP207, and AP193)

**Figure 10:**
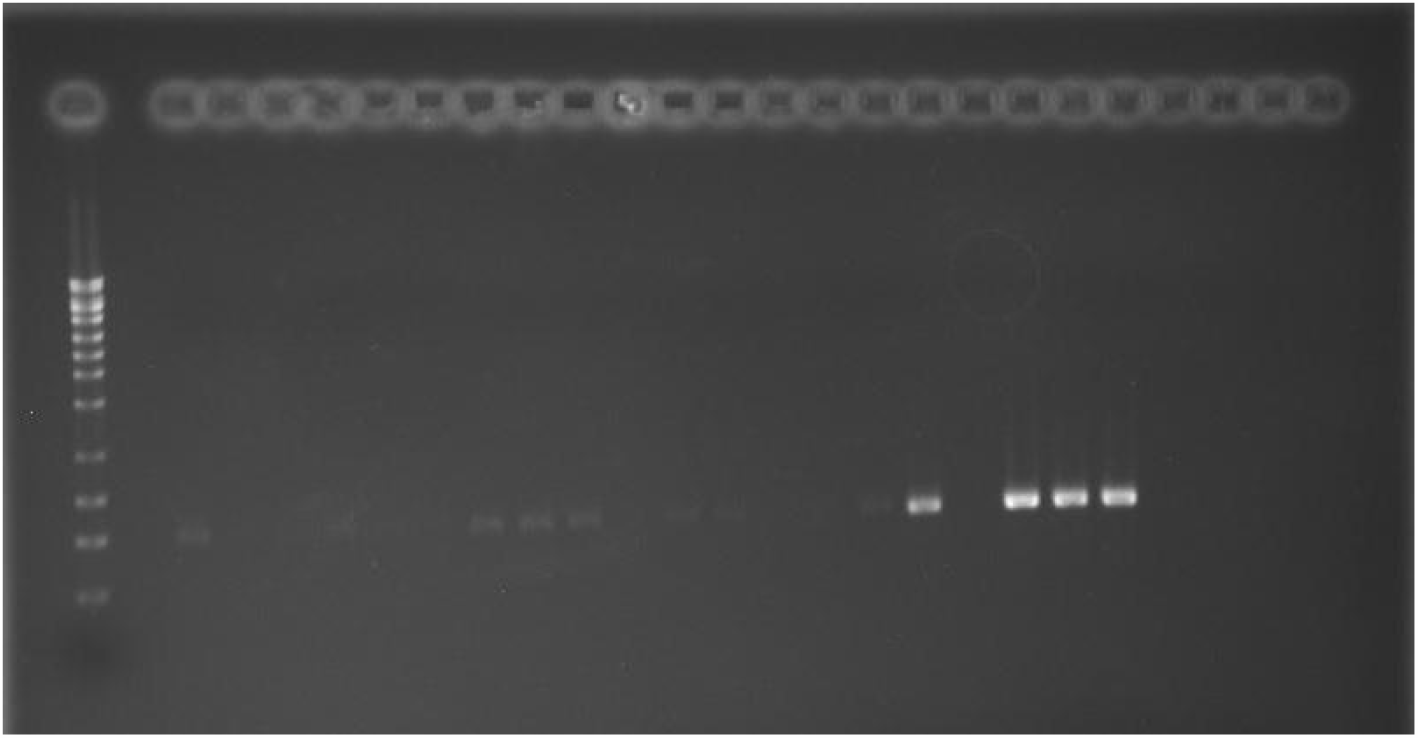
*exuT* gene detected in Bap strains (AP211, AP219, AP296, AP75, AP299, AP205, AP301, AP216, AP143, AP150, AP202, AP241, and AP193)

**Figure 11:**
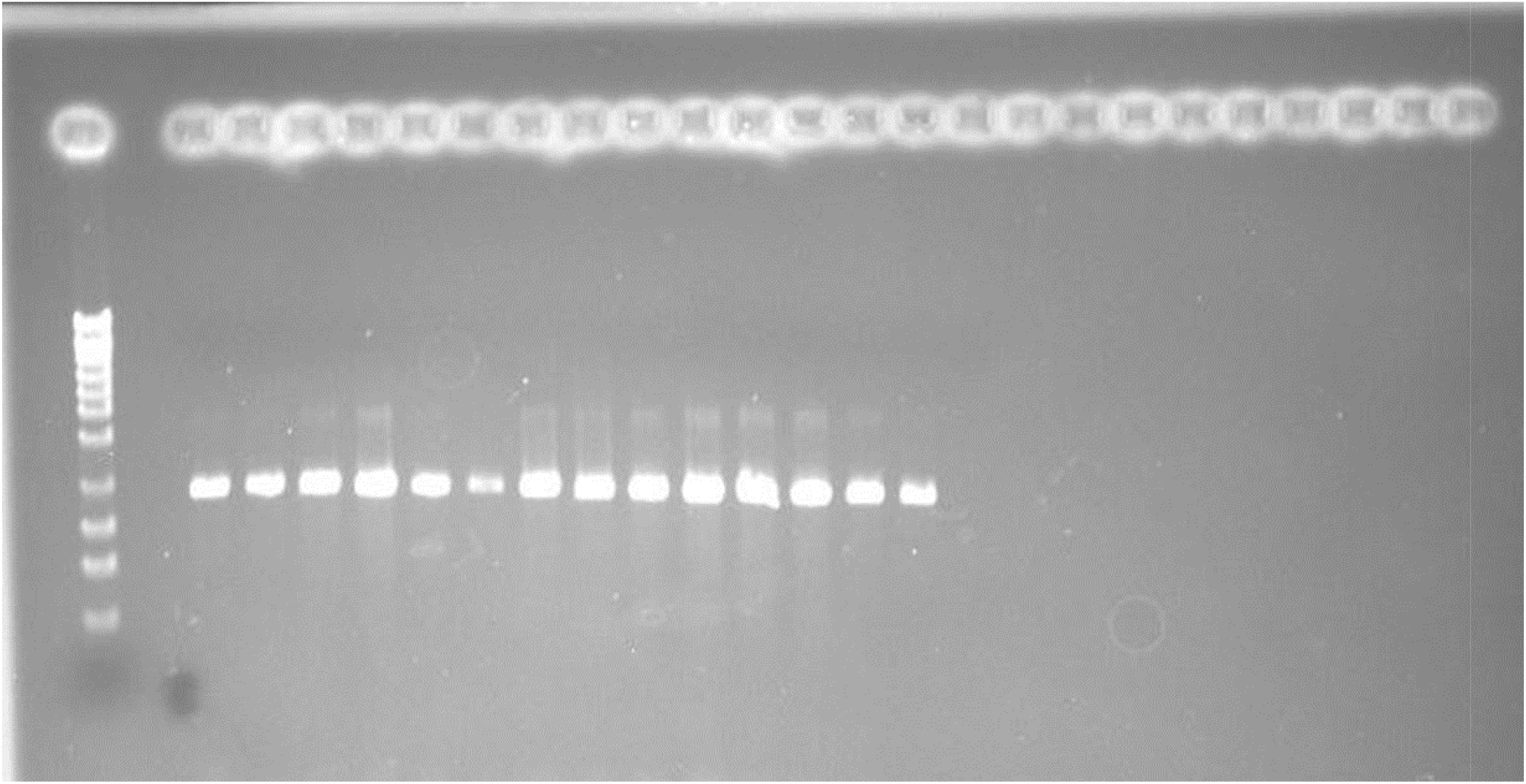
*uxuB* gene detecte din Baps trains(A P2 97,AP80, AP.86, and AP193)

**Figure 12:**
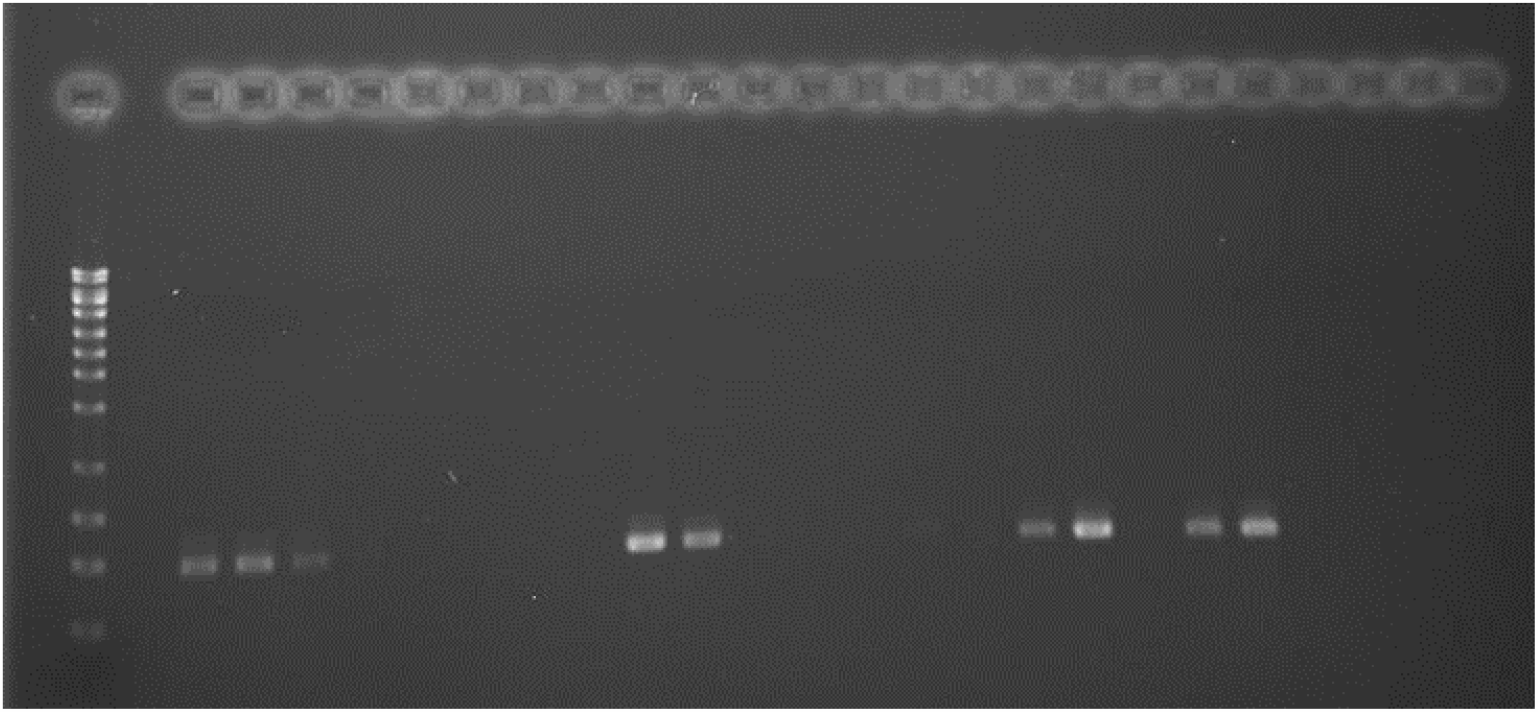
*uxuB* gene detected in Bap strains (AP211, AP295, AP205, AP241, AP184, AP207, and AP193)

**Figure 13:**
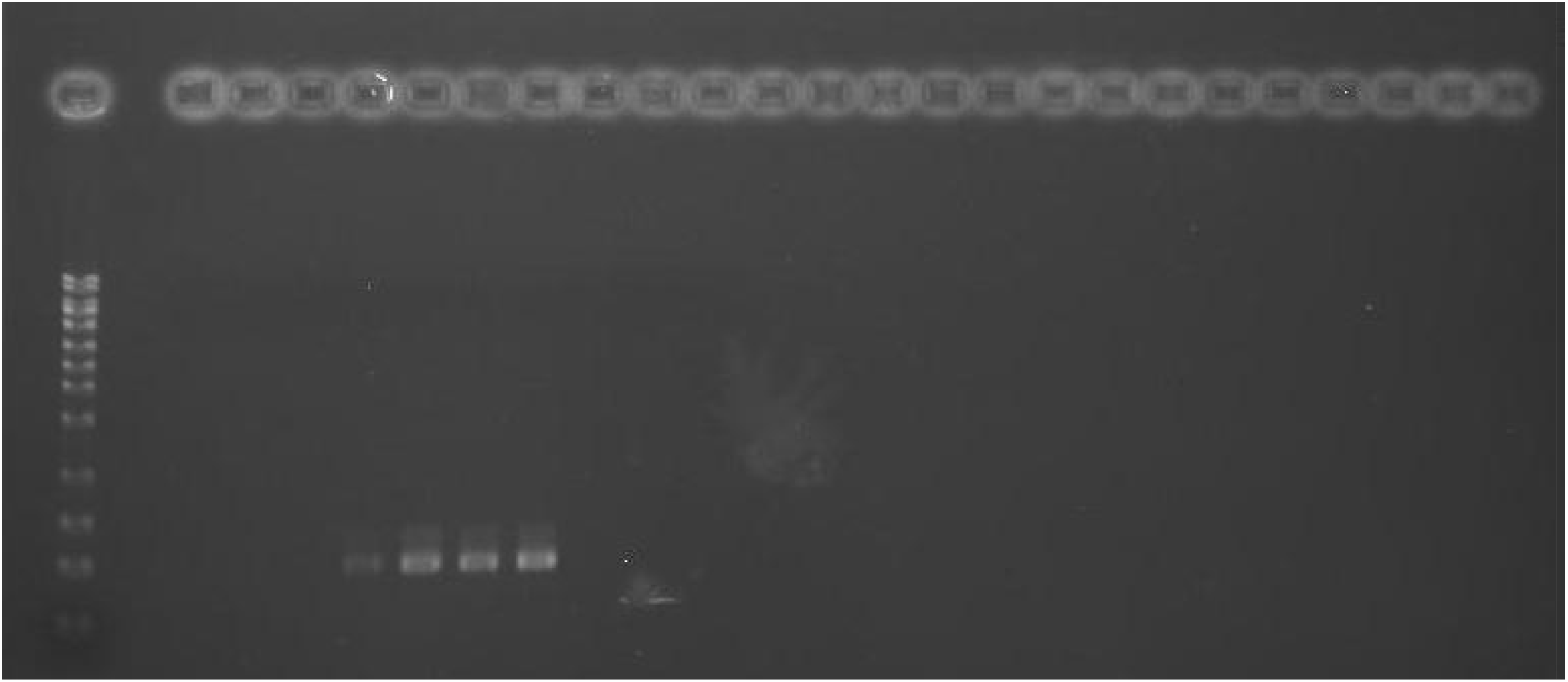
*uxuB* gene detected in Bap strains (AP296, AP241, and AP193)

**Figure 14:**
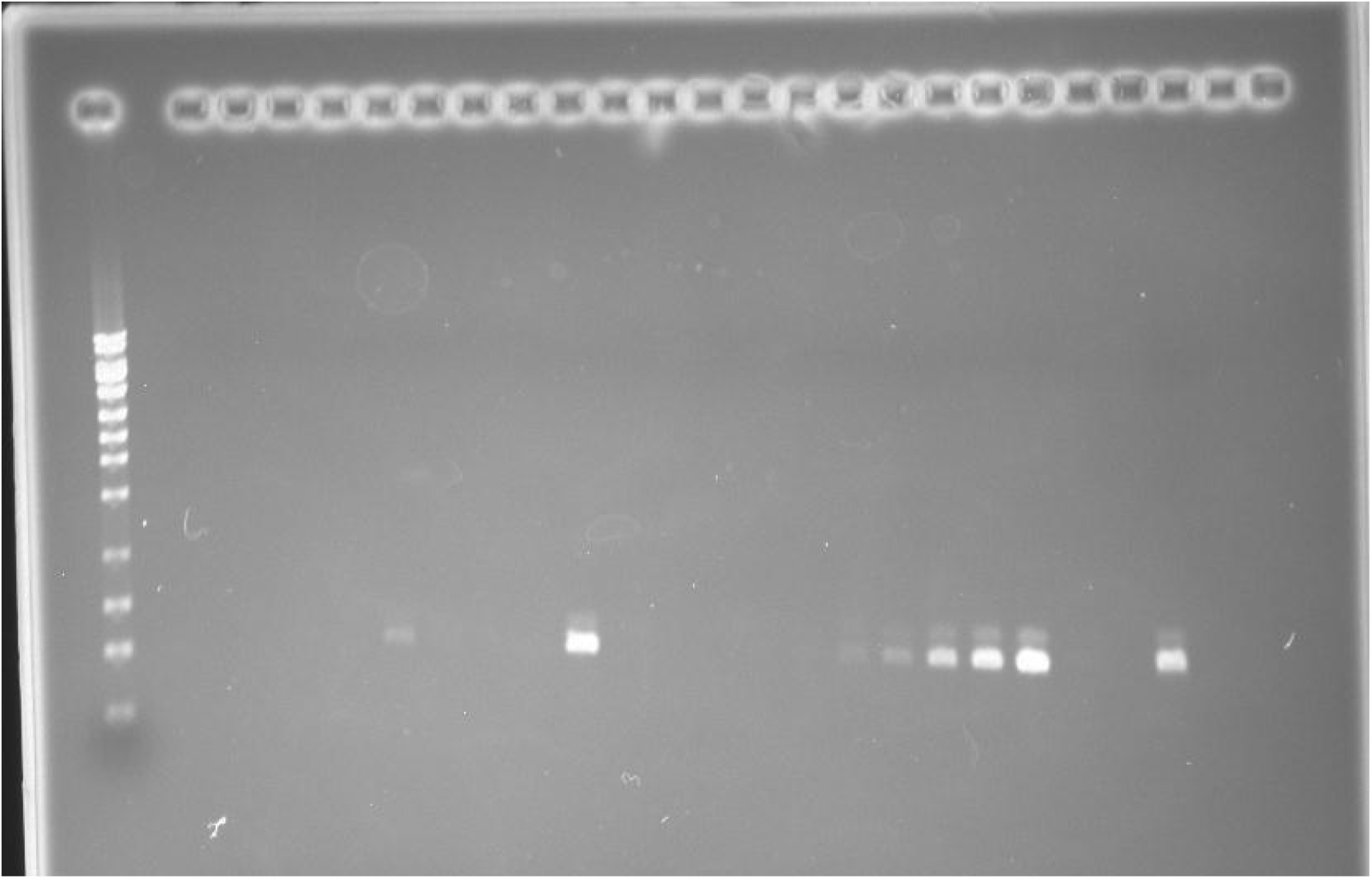
*uxuB* gene detected in Bap strains (AP215, AP108, AP188, AP213, and AP193)

**Figure 15:**
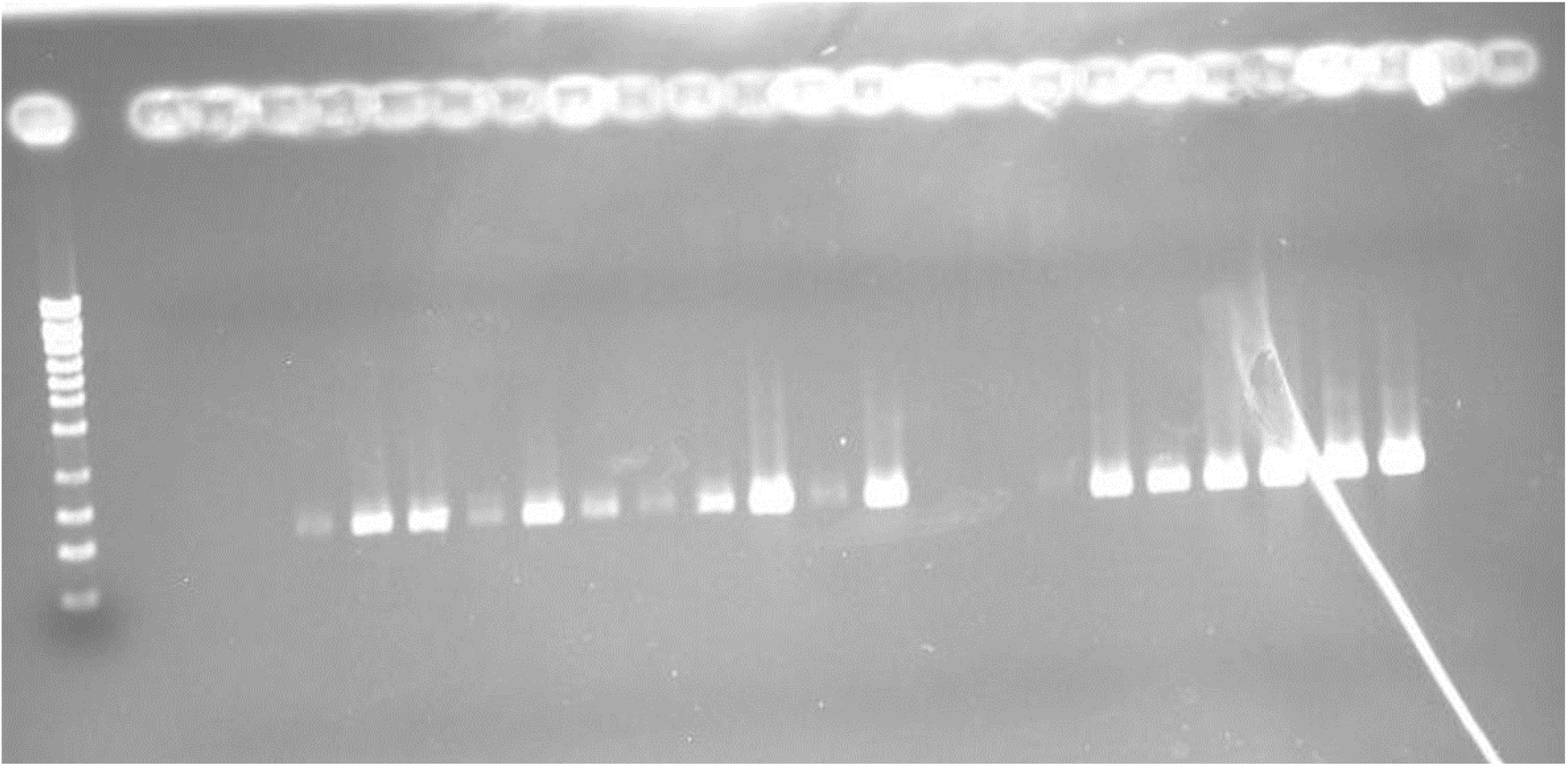
*uxuB* gene detected in Bap strains (AP201, AP71, AP200, AP87, AP183, AP136, AP203, AP75, AP196, AP78, AP305, AP299, and AP193)

**Figure 16:**
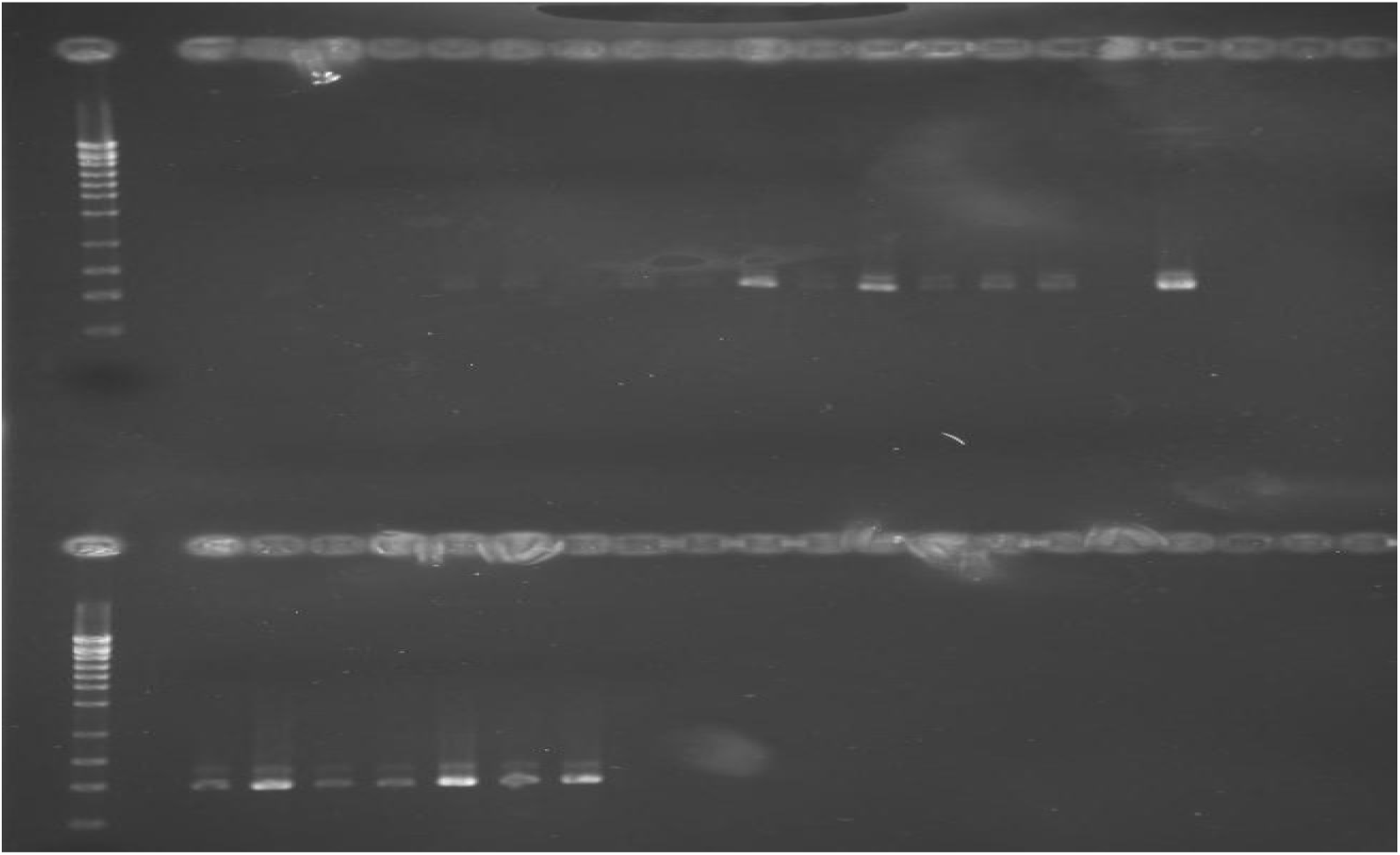
*uxuB* gene detected in Bap strains (AP216, AP150, AP190, AP81, AP208, and AP193)

**Figure 17:**
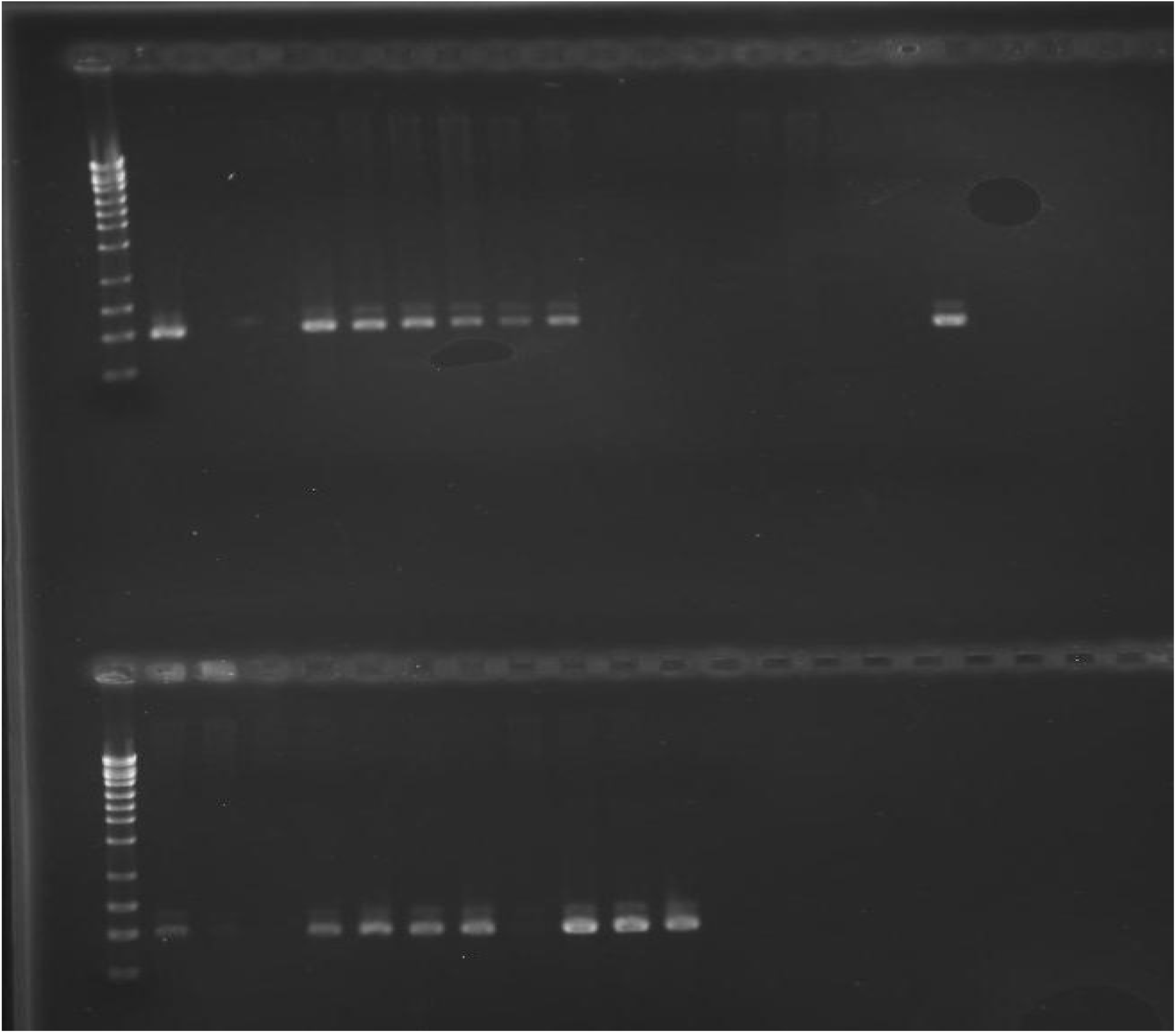
*uxuB* gene detected in Bap strains (AP191, AP195, AP212, AP210, AP79, AP77, AP189, AP67, AP300, AP260, AP304, AP52, AP135, AP76, and AP193)

**Figure 18:**
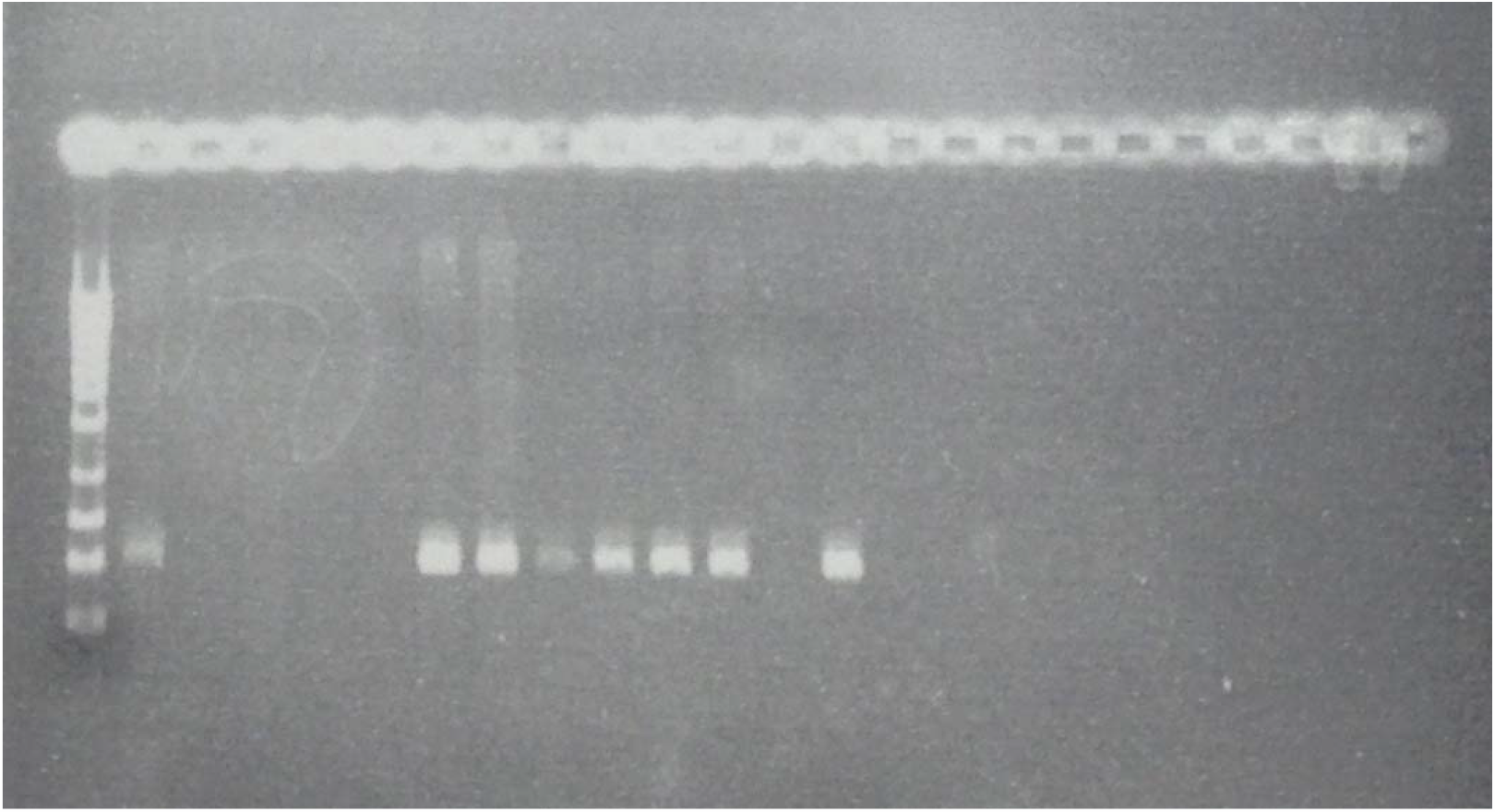
*uxuB* gene detected in Bap strains (AP198, AP192, AP298, AP194, AP85, AP112, and AP193)

**Figure 19:**
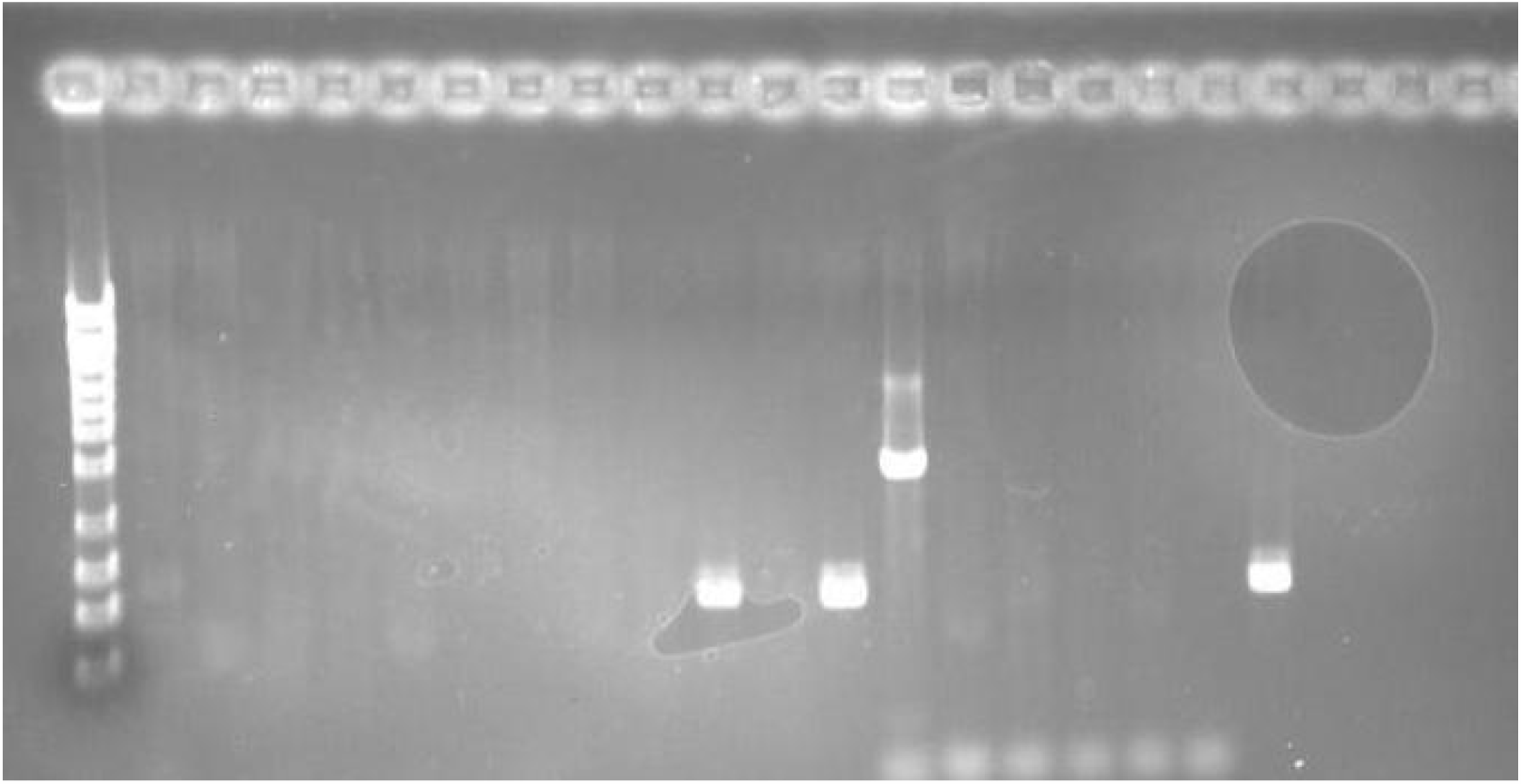
*uxuB* gene detected in Bap strains (AP143, AP197, and AP193)

**Figure 20:**
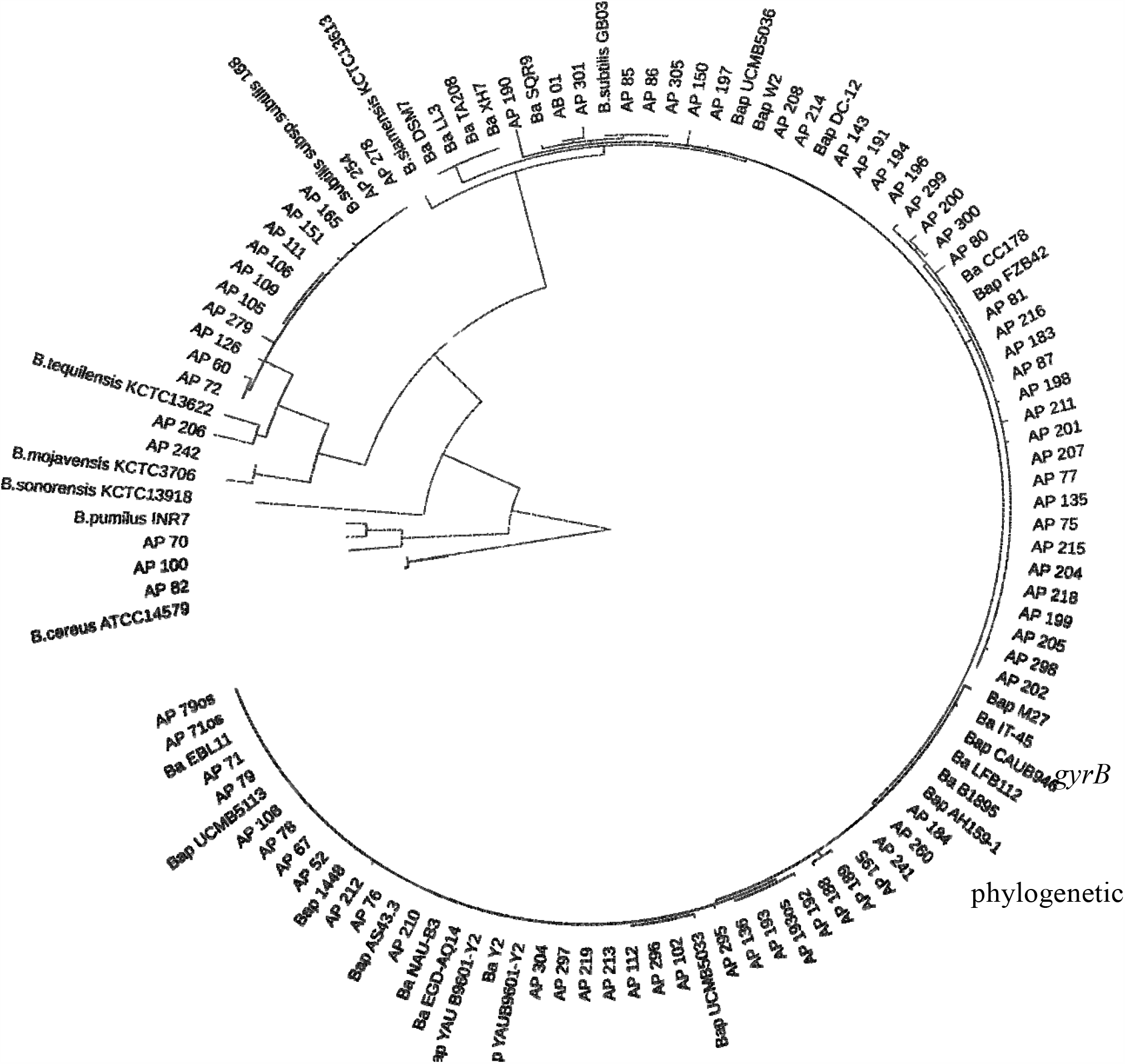
*gyrB* phylogenetic tree of 79 Bacillus spp.

